# Dynamics of aqueous suspension of short Hyaluronic acid chains near DPPC bilayer

**DOI:** 10.1101/2024.02.14.580078

**Authors:** Anirban Paul, Jaydeb Chakrabarti

## Abstract

The synergy between Hyaluronic acid (HA) and lipid molecules plays a crucial role in synovial fluids, cell coatings, etc. Diseased cells in cancer and arthritis, show changes in HA concentration and chain size, impacting the viscoelastic and mechanical properties of the cells. Although the solution behavior of HA is known in experiments, a molecular-level understanding of the role of HA on the dynamics at the interface of HA-water and the cellular boundary is lacking. Here we perform atomistic molecular dynamics simulation of short HA chains in explicit water solvent in presence of DPPC bilayer, relevant in pathological cases. We identify a stable interface between HA-water and the bilayer where the water molecules are in contact with the bilayer and the HA chains are located away without any direct contact. Both translation and rotation of the interfacial waters in contact with the lipid bilayer and translation of the HA chains exhibit subdiffusive behavior. The diffusive behavior sets in slightly away from the bilayer, where the diffusion coefficients of water and HA decrease monotonically with increase in HA concentration. On the contrary, the dependence on HA chain size is only marginal due to enhanced chain flexibility as their size increases.

**TOC Graphic:** 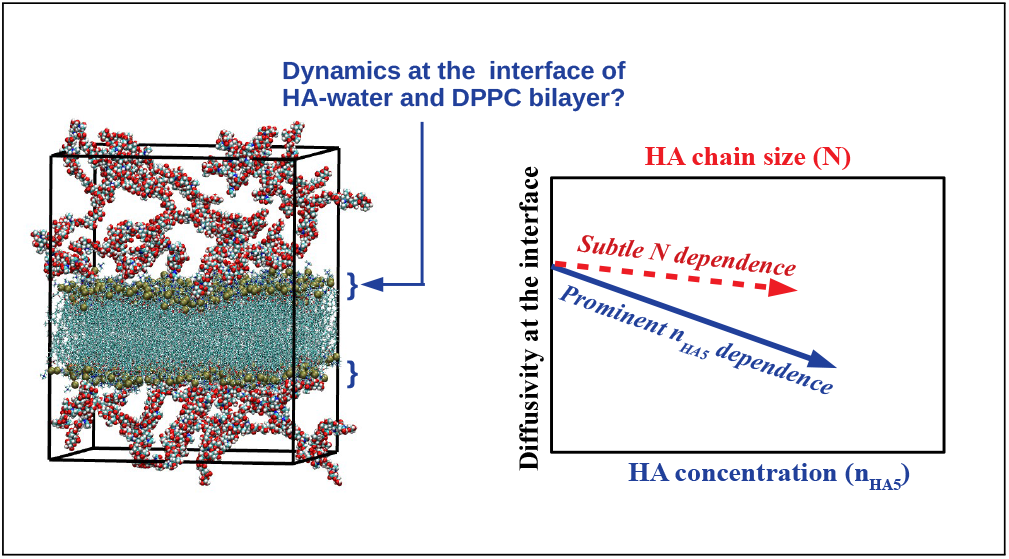

---

Hyaluronic Acid (HA) is a hydrophilic, polyanionic glycosaminoglycan molecule with molecular weight ranging from 10 kDa to 10,000 kDa^1–3^. It is an essential component of tissues and bodily fluids. The organization of HA chains in the vicinity of cell membranes and the synergy between HA and lipid molecules play vital roles in biology, such as maintaining the viscoelasticity of the synovial fluids, elasticity of cells and so on ^4–7^. The abundance of low molecular weight HA is reported in many studies in pathological conditions like cancer^1,8,9^, arthritis^10^, etc. Although the bulk properties of HA solutions have been widely explored ^11–13^, there has been very limited understanding of the dynamics at the interface of the aqueous solution of short HA chains in the vicinity of cell membranes.

HA provides extreme lubrication to synovial joints along with lubricin and phosphatidyl-choline^14^. The degradation of HA chains in the synovial fluid causes inflammation and arthritis of the joints. The low molecular weight HA chains (< 200 KDa) are more prevalent in the vicinity of lipid vesicles in cancer cells as well, compared to normal cells ^9,15^. The low molecular weight short HA chains are potential biomarkers for early detection of colon cancer cells^8^. Experiments demonstrate that the alternation of HA content leads to mechanical changes in diseased cells such as in the inflamed pancreatic tissues^16^. Short HA-coated extra-cellular vesicles (EVs) secreted from colon cancer cells are also observed to be significantly more elastic than the normal cell-derived EVs ^17^. The observations have been explained in terms of the changes in water ordering in presence of short HA chains using Molecular Dynamics simulations. It has also been reported that the water molecules are slower near the cancer vesicles when the HAs around the vesicles are degraded using the hyaluronidase enzyme^17^. However, due to the presence of hyaluronidase and the complex environment of the cancer cell in vivo, the effect of HA concentration and HA chain size on the dynamics of the water molecules cannot be decisively inferred. Recently Zhang et al. have also shown using molecular dynamics simulation that the short HA chains affect the water mobility and permeation rate of water through Aquaporin proteins ^18^. Nonetheless, to the best of our knowledge, there has been no attempt so far for a detailed understanding of the dynamical nature of the micro-environment near cell membranes in presence of short HA chains, despite their importance in many contexts including therapeutic development.^19,20^

With this backdrop, we perform in-silico Molecular Dynamics (MD) simulations on an atomistic model system, consisting of an aqueous solution of short HA chains in the vicinity of a Dipalmitoylphosphatidylcholine (DPPC) bilayer. Our system models an extra-cellular medium in the vicinity of the cancer cell membrane which is typically abundant in phosphatidylcholine lipids^21,22^and shorter HA chains^8,9^. We investigate the effect of HA monomer concentration and HA chain length, in the low molecular weight (mw) limit, on the interfacial dynamics near the lipid bilayer. We characterize the dynamics in terms of the residence time of water molecules^23^ and HA chains in the vicinity of the bilayer, their translational and rotational mean squared displacements^24–26^, and the diffusion coefficients.

We observe that the water molecules and HA chains form a stable interface with the lipid bilayer where water comes in direct contact with the bilayer, while the HA chains stay slightly away from the bilayer without direct contact. The residence times of the water molecules at the HA-water and DPPC interface are around tens of pico-seconds (ps), while those of HA monomers are around hundreds of pico-seconds. The residence times increase with HA concentration and HA molecular length. We further observe that the dynamical behaviors at the interface respond differently to HA monomer concentration and HA chain size. The contact layer of water with the bilayer shows sub-diffusive dynamics both in translation parallel to the bilayer plane and rotation. The respective mean squared displacements (MSD) of these molecules increase with time with exponents less than unity. The diffusive motion of the water molecules, where the MSDs increase linearly in time, sets in beyond the contact layer. In this region, both translational and rotational diffusion coefficients decrease as HA concentration but not sensitive to the chain size. The HA chains exhibit sub-diffusive translation but diffusive rotational motion at the interface. The rotational diffusion coefficient of the chains decreases with increasing HA concentration but this quantity does not depend on HA chain sizes for large chains. In addition, we find that the in-plane translational diffusion of the lipid headgroups gets enhanced with increasing HA concentration, but varies marginally with HA chain size. On the other hand, the rotational diffusion of lipid PN vectors decreases with HA concentration but increases monotonically with HA chain size.

## Results

In this report, we consider three distinct molecular sizes of HA. HA monomers are denoted as HA1 (mw = 0.3 kDa), pentamers are denoted as HA5 (mw = 1.9 kDa), and decamers are described as HA10 (mw = 3.8 kDa). We consider systems comprising 10, 30, and 50 HA5 chains (n_HA5_ = 10, 30, and 50) to observe the effect of HA concentration and systems with different HA chain sizes for a fixed number (150) of HA monomers, namely, 150 HA1 monomer (N=1), 30 HA5 pentamer (N=5), and 15 HA10 decamer (N=10) to investigate the effect of HA chain size. Equilibrium snapshots of all the cases are shown in SI Figure S1, which are similar to those reported in the previous study^17^. The equilibration in each case is confirmed by the convergence of the area per lipid of the DPPC bilayer as done in our previous work^17^. The interface between HA-water and DPPC phases is computed from the equilibrium density profiles of the molecular constituents. Their mean residence time at the interface is estimated from the survival probability data. Furthermore, we compute the translational and rotational mean squared displacements (MSD) of water, HA chains while they reside at the interface and the lipid headgroups. The diffusion coefficients are acquired from the slope of MSD data at the long time limit if the data show linear time dependence, namely, diffusive region.

Before going into the detailed dynamical properties of the ternary system (HA, water, and DPPC) at the interface, we determine the location of the HA-water interface with the DPPC bilayer from the equilibrium density profiles of the molecular constituents as reported earlier^17^. The equilibrium density profiles of the phosphorus atoms of the lipid bilayer (*ρ*_*P*_ (z)), oxygen atoms of the water molecules (*ρ*_*W*_ (z)), and the center of mass (COM) of the HA monomers (*ρ*_*H*_ (z)) are computed along the bilayer normal direction (z-axis) with respect to the origin at the center of the bilayer. *ρ*_*P*_ (z), *ρ*_*W*_ (z) and *ρ*_*H*_ (z) for two cases n_HA5_=30 and N=10 are shown here in Figure 1(a) and 1(b) respectively. Density profiles for other cases are shown in SI Figure S2. The peaks of *ρ*_*P*_ (z) are at *d*_*P*_ ^*l*^ is at -20 Å and *d*_*P*_ ^*u*^ at 20 Å, corresponding to the phosphorus atoms of the two leaflets. Thus, the bilayer thickness |*d*_*P*_ ^*u*^ − *d*_*P*_ ^*l*^| = 40 Å is in agreement with the previous report^27^.The water density profile *ρ*_*W*_ (z) reaches its bulk value at z=± 35 Å from the bilayer center. Any chemical species of the system is considered to be at the interface if its z coordinate satisfies the following condition: either *d*_*P*_ ^*l*^ − 15 Å < z < *d*_*P*_ ^*l*^ or *d*_*P*_ ^*u*^ < z < *d*_*P*_ ^*u*^ + 15 Å. We observe that the interface width (∼ 15 Å) is not so sensitive to n_HA5_ and N. We also note that the peak of *ρ*_*H*_ (z) is located at almost where the bulk density of water sets in and *ρ*_*H*_ (z) is very low till 5 Å distance from the phosphate peaks showing that HA forms a non-contact interface with DPPC in agreement to the previous studies^17,28^.

**Figure 1.**
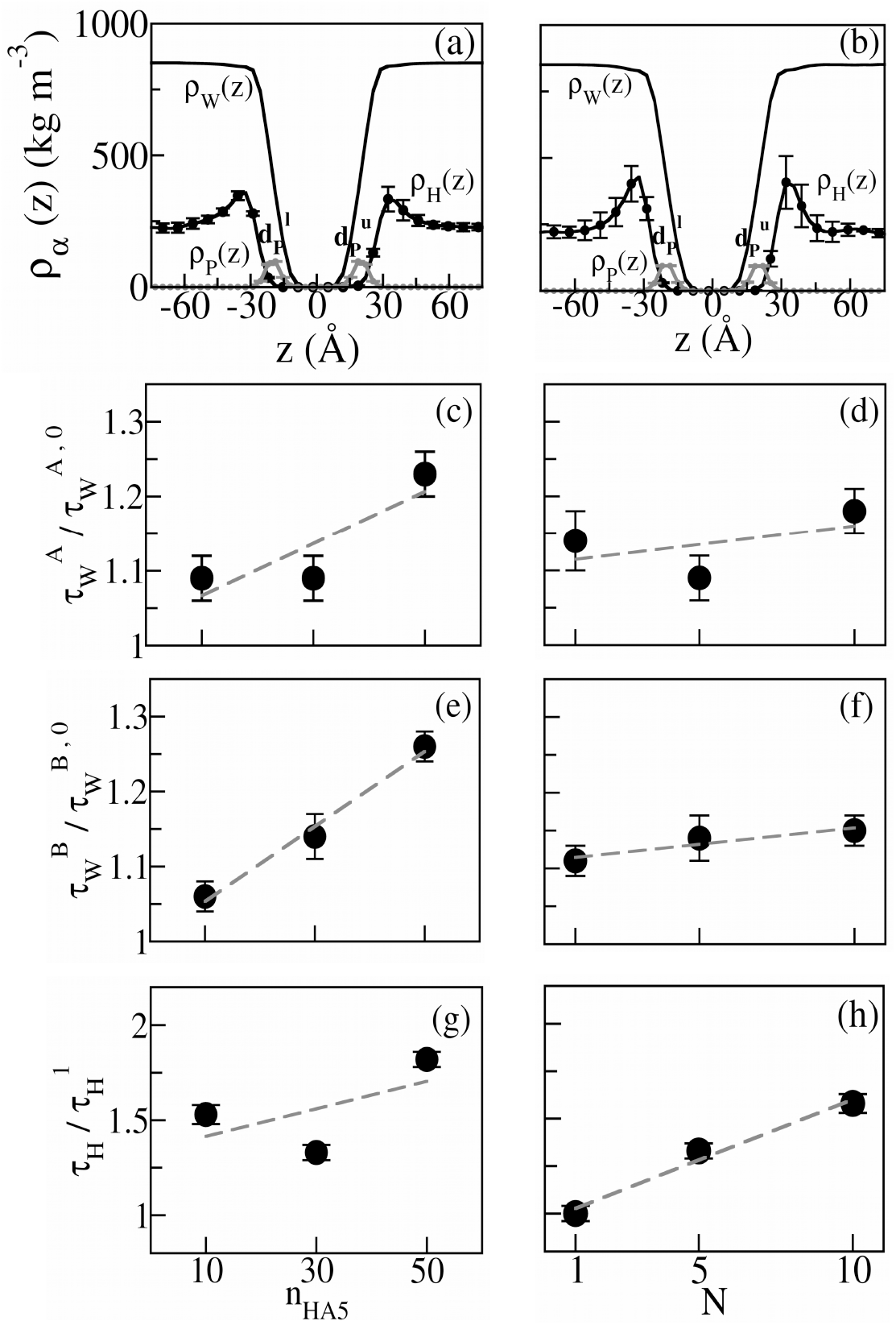
(a) Density profiles of phosphorus atoms (*ρ*_*P*_ (z), *grey solid line*), HA (*ρ*_*H*_(z), *dotted black line*), and water (*ρ*_*W*_ (z), *solid black line*) along the bilayer normal for n_HA5_=30 and (b) for N=10. Origin is set at the bilayer center. *d*_*P*_ ^*l*^ and *d*_*P*_ ^*u*^ show the peaks of *ρ*_*P*_ corresponding to the phosphorus atoms of the lower and upper leaflets of the bilayer. *ρ*_*H*_(z) is amplified by 5 times for better clarity. (c) The Mean residence time of water molecules in region 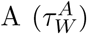 for varying n_HA5_ and (d) for different N. (e) Mean residence time of water molecules in region 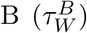 for varying n_HA5_ and (f) for different N. The respective residence times are scaled with the quantities for HA-free cases: 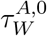 and 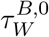. (g) The mean residence time of HA chains in region 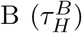 for different n_HA5_ and (h) for different N. The data are scaled with 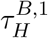, the mean residence time of HA chains in region B. The best linear fits are indicated by the *dashed gray* lines.

### Residence time of water and HA chains at the interface

We consider two distinct regions in the interface while probing the dynamics. Region A extending up to 5 Å below *d*_*p*_^*l*^ or above *d*_*p*_^*u*^ where only the water molecules reside; and Region B extending from 5 Å to 15 Å from the *d*_*p*_^*l*^ or *d*_*p*_^*u*^ consisting of both water and HA chains. We compute the survival probabilities of water S_*W*_ (t) in these regions using the coordinates of the oxygen atoms (SI Figure S3). The mean residence times of the interfacial waters, 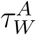 and 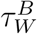 are obtained from S_*W*_ (t) data in respective regions(SI table S1). 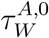 and 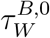 indicate the residence time of water molecules in region A and B respectively for HA free case (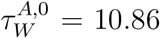 ps and 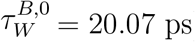). Here we find 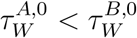, for region B has a larger width than A. 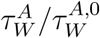 data for different n_HA5_ and N are shown in Figure 1(c) and 1(d) respectively. Figure 1(e) and 1(f) shows 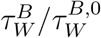 data for different n_HA5_ and N respectively. In both regions, water molecules get slower in presence of HA molecules than in the HA-free case. Both 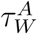 and 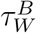 decrease with n_HA5_ but remain almost the same with N. The residence time of the HA chains, *τ*_*H*_, is computed from the survival probability data (S_*H*_ (t)) of COM of at least one monomer of a chain within the stipulated interfacial region (SI Figure S4). We find that *τ*_*H*_ is an order of magnitude slower than the water molecules (See SI table S2). Figure 1(g) and 1(h) show 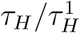 data for different n_HA5_ and N respectively. Here 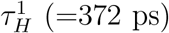 implies HA monomer residence time for N=1. Our data suggest that *τ*_*H*_ increases linearly with n_HA5_ and N. But the change with N is less pronounced. Thus both water and HA chains spend a longer time in the interface with increasing n_HA5_, while the time does not change much with N.

### Dynamics of interfacial water molecules

Next, we study the dynamics of water at the interface of HA and DPPC bilayer. We compute two-dimensional translational mean squared displacements (MSD), 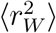 in the plane parallel to the bilayer and rotational MSD, 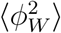 of the water molecules for both region A and region B. We make sure that the calculations of 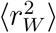 and 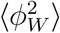 are performed while the oxygen atoms of the waters reside continuously within a given region^23^. 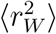 and 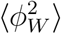 evolves with time with exponents *β*_*W*_ and *γ*_*W*_ respectively i.e 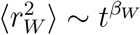 and 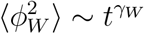. 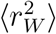 in region A for different n_HA5_ and N are shown in SI Figure S5. We fit the log-log data of 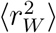 versus Δ*t* up to water residence time in region A, 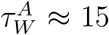 ps to compute the translational exponent *β*_*W*_. We find that *β*_*W*_ in region A is close to 0.75 for all HA concentration and chain size ranges, indicating sub-diffusive motion of water molecules (SI Table S3), as observed in previous studies^29^.

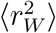 in region B for different n_HA5_ and N are shown in Figure 2(a) and 2(b) respectively. In this region, using 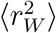 data up to water residence time 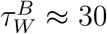 ps we find that the *β*_*W*_ values are close to unity, suggesting diffusive dynamics. We fit 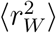 data with 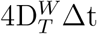 to extract the two-dimensional diffusion coefficient 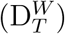 of the interfacial waters in the plane parallel to the bilayer^30^. In the absence of HA, the diffusion coefficient of the interfacial water molecules in region B, D^W,0^_T_ = 6.19 ± 0.07 × 10 ^*−*5^ cm^2^/s. The diffusion coefficients of the interfacial water molecules scaled to the HA free case, D^W^_T_/D^W,0^_T_ are shown in Figure 2(c) and 2(d) as functions of n_HA5_ and N. We observe that D^W^_T_ for all cases are smaller than D^W,0^_T_, implying that HA restricts the translation of the interfacial waters. D^W^_T_/D^W,0^_T_ decreases linearly with n_HA5_. On the other hand, D^W^_T_/D^W,0^_T_ in Figure 2(d) shows a marginal linear decrease with N.

**Figure 2.**
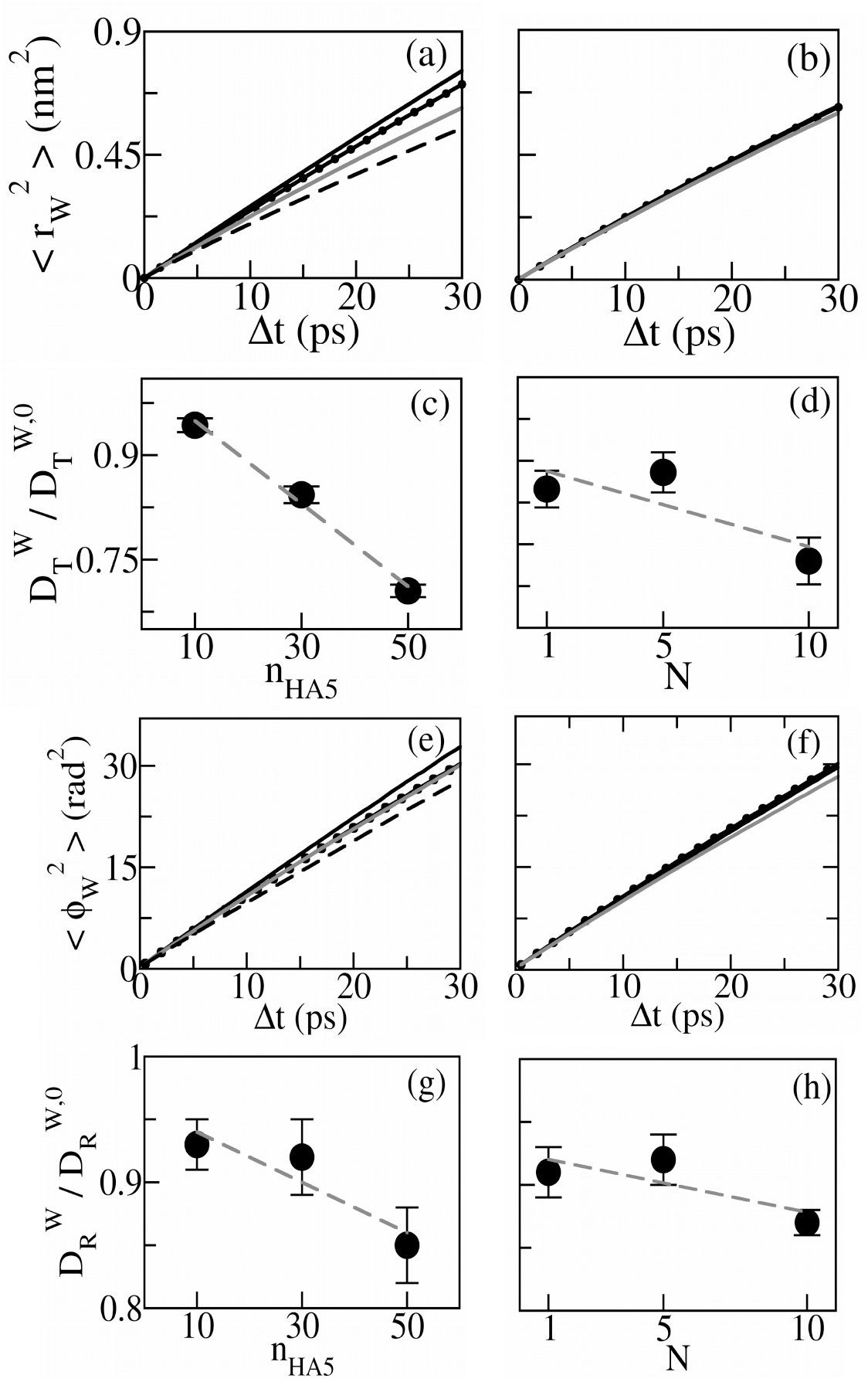
(a) Translational MSD of the water molecules 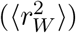 in region B, in the plane parallel to the bilayer surface, for different HA concentrations: n_HA5_=0 (*solid black line*), 10 (*dotted black line*), 30 (*solid gray line*) and 50 (*dashed black line*) (b) 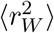 in region B for varying HA chain sizes: N=1 (*solid black line*), N=5 (*dotted black line*) and N=10 (*solid gray line*). (c) Two-dimensional diffusion coefficients of the water molecules 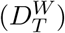 in the diffusive region (region B) for different n_HA5_ and (d) N. The values are scaled by 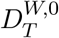, the two-dimensional diffusion coefficient of water for HA free case. The best linear fits are shown by the *dashed gray* lines. (e) Rotational MSD of water molecules 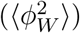 in region B for varying n_HA5_ and (f) for different N. The same line types as (a) and (b) are used. (g) Rotational diffusion coefficients of water molecules 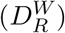 in region B for different n_HA5_ and (h) N. The values are scaled by rotational diffusion coefficient 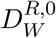 for HA free case. The *dashed gray* lines indicate the best linear fits.

We also investigate the rotational motion of the water molecules in regions A and B. We use the vector rotational displacements 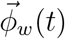 of the water dipole vectors 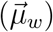 in time t to compute the rotational MSD, 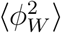^24,31^ (see details in Methods). 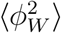 of the interfacial water molecules, in region A for varying n_HA5_ and N are shown in SI Figure S6. Fitting the log-log data of 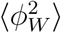 with a power function up to 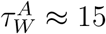 ps, we find that the corresponding exponent of the temporal dependence *γ*_*W*_ is close to 0.73 (SI Table S4), which suggest sub-diffusive rotational motion of water in region A^29^.

For region B, we show 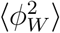 in Figure 2(e) and 2(f) for different HA concentrations and HA chain sizes. Using 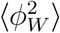 data up to 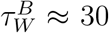, we observe diffusive rotational motion with the exponent *γ*_*W*_ close to unity, as the translational counterpart. In the linear time dependence region (region B), the rotational diffusion coefficient is extracted from the slope of the 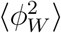 vs Δ*t* data^24,32^as done for translational motion. We find the rotational diffusion coefficients of the interfacial waters in region B for HA free case, D^W,0^_R_ = 0.27rad^2^/ps. We show in Figure 2(g) and Figure 2(h) 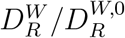 data as functions of n_HA5_ and N respectively.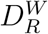 in all cases are smaller than 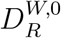, suggesting that HA makes the interfacial water molecules rotationally slower. We find a linear increase in both cases implying slower rotational diffusion, although the N dependence is less pronounced than the n_HA5_ dependence as noted in the translational diffusion as well.

Experimental techniques such as infrared absorption, Raman scattering, and NMR, report rotational correlation times of water, 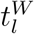, where 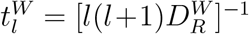 and l corresponding to the l’th Legendre polynomial^33^. We compute the first rank rotational correlation time 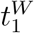 from the rotational autocorrelation function C(*t*) of the water molecules^23^ in region B (SI Figure S7) for different HA concentrations and HA chain sizes (SI table S5). We find that the rotational auto-correlation time for water increases in presence of HA as observed experimentally^11,12^. We observe that 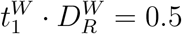 that corresponds to l=1.

### Dynamics of the interfacial HA chains

Next, we investigate the dynamics of the HA chains at the HA-water and DPPC interface. We compute the translational MSD of the HA chains, 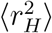 in the plane parallel to the bilayer surface and the rotational MSD of the HA chains 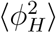 at the interface. To apprehend the interfacial dynamics, 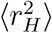 is calculated for only those chains which have at least one monomer present at the interface and using the positions of those interfacial monomers till their residence time. Whereas, 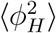 are calculated using the vector rotational displacement 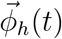 of the HA end-to-end vector in time t (see details in Methods) provided that at least one end monomer of the chain belongs to the interface. For N=1, we use the long molecular axis of the interfacial HA monomers to compute rotational MSD.

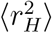 vs Δ*t* for different n_HA5_ and N are shown in Figure 3(a) and Figure 3(b) respectively. The exponents *β*_*H*_ of temporal dependencies of 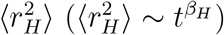 are obtained from the log-log data of 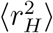 vs time Δt up to *τ*_*H*_ ≈ 500ps. We find that 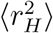 exhibit sub-diffusive behavior within HA residence at the interface, with *β*_*H*_ in the range of 0.80 to 0.88 (SI Table S6). This observation is similar to the subdiffusion of polyanionic DNA chains adsorbed to lipid bilayer at a short timescale^34,35^. Figure 3(c) and 3(d) describe 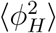 vs Δt plot for different n_HA5_ and N respectively. We observe that the exponents of temporal dependence of 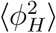 upto *τ*_*H*_ ≈ 500ps is close to unity, suggesting that the HA rotational motion is diffusive. The rotational diffusion constants 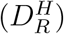 are computed from the slope of the time dependent 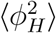 data. For HA monomers 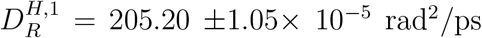. 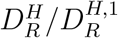 data are shown in Figure 3(e) for different n_HA5_. We note that 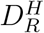 for any n_HA5_ undergoes an order of magnitude decrease compared to 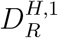 and further decreases linearly with increasing n_HA5_. Figure 3(f) describes 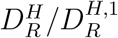 data for varying N. 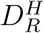 for N=5 and N=10 are dramatically less than 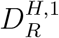 (almost 100 times) but no significant change from N=5 to N=10. In addition, we estimate the first rank rotational auto-correlation time 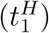 of the HA chains from 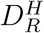 data for *l* = 1 (SI Table S7). We observe that 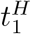 increases as n_HA5_ and N increases, but less sensitive with N for its higher values. We also note that 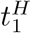 are of the order of nanoseconds, where 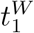 of water are in the picosecond range.

**Figure 3.**
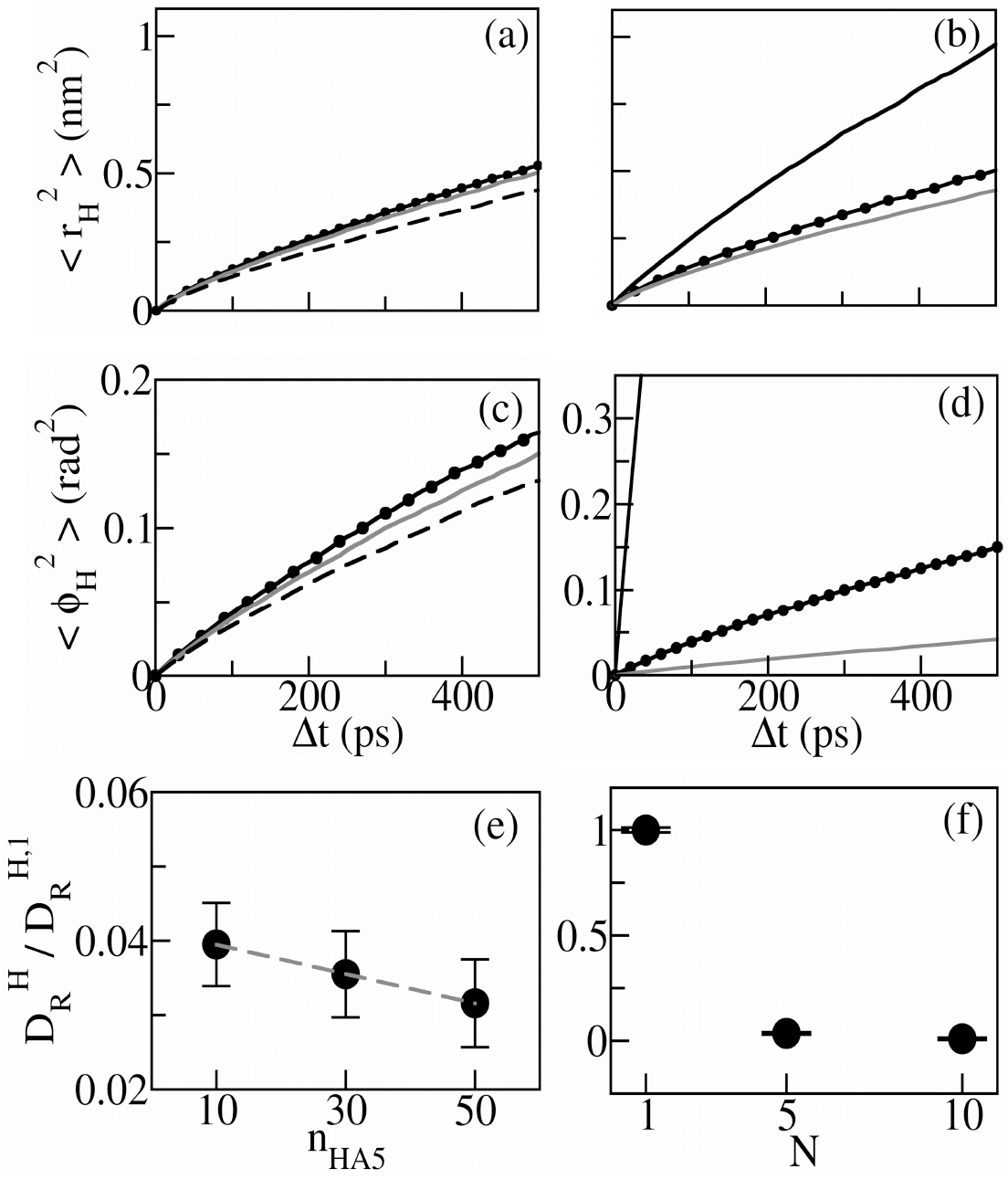
(a) Translational MSD of HA chains, 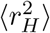, in the plane parallel to the bilayer surface, for n_HA5_=10 (*dotted black line*), n_HA5_=30 (*solid gray line*) and n_HA5_=50 (*broken black line*) and for N=1 (*solid black line*), N=5 (*dotted black line*) and N=10 (*solid gray line*). (c) Rotational MSD of HA chains, 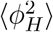 for different n_HA5_ and (d) for different N. Same line types as (a) and (b) are used. (e) Rotational diffusion coefficients, 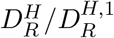 data for different n_HA5_ and (f) for different N. The broken line shows the best-fitted straight line. Error bars are smaller than the symbol size.

### Dynamics of lipid head groups

The influences of HA on the lipid headgroup dynamics are investigated by computing the translational MSD of the phosphorus atoms^36,37^and rotational MSD of the lipid headgroup PN vectors^26^. The translational MSD 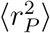 is computed in the lateral plane of the bilayer surface. SI Figure S8 shows 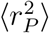 for different HA concentrations and HA chain sizes. We observe an initial sub-diffusion in 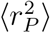 in the time range Δ*t* < 50 ns, followed by the diffusive regime, consistent with the earlier studies ^36,38^.

The in-plane translational diffusion coefficients of the phosphorus atoms 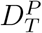 are obtained from the slope of 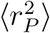 versus Δ*t* data in the diffusive region. For HA free case, we find the phosphorus diffusion, 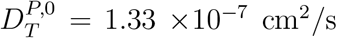. We describe 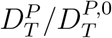 data for different n_HA5_ in Figure 4(a). We note that the lipid phosphorus diffusion increases with n_HA5_ and exceeds 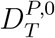 at large n_HA5_. This is quite the opposite to the reduction of lipid diffusion at high sucrose and trehalose concentrations^39–41^. The higher mobility of the phosphorus atoms in presence of HA is consistent with the bilayer flexibility observed previously^17^. The N dependence of 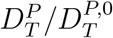 in Figure 4(b) is weaker showing a maximum around N=5.

**Figure 4.**
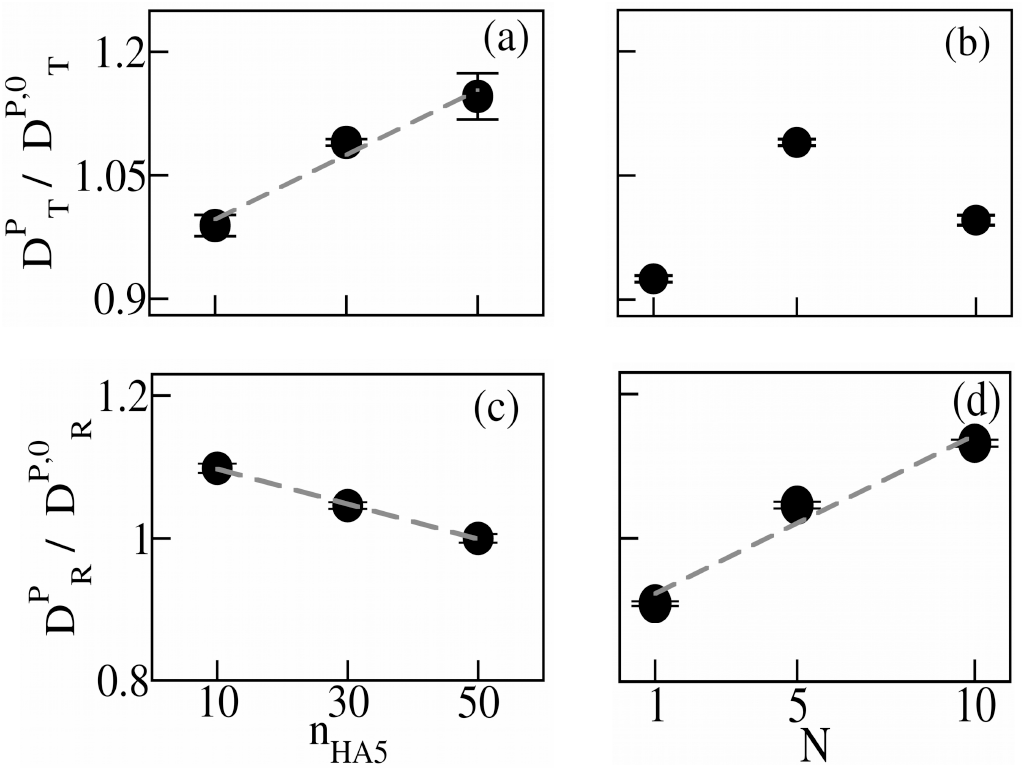
(a) Lateral diffusion coefficient of the phosphorus atoms of the lipid bilayer 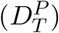 for different HA concentrations and (b) for different HA chain sizes. (c) The rotational diffusion coefficients of the lipid PN vectors 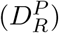 for varying HA concentrations and (d) for different chain sizes. 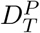 and 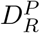 are scaled with the diffusion coefficients 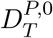 and 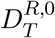 respectively for HA free case. Best linear fits are shown by the *dashed gray* lines. Error bars are smaller than the symbol size.

Rotational MSD of the PN vectors of the lipid molecules, 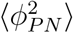 are computed with the unbounded variable 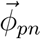 using their projection vectors on the x-y plane^26^. 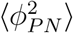 for different n_HA5_ and N are shown in SI Figure S9. The exponent of temporal dependence of 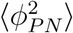 is close to unity in all the cases so that one can extract the rotational diffusion coefficient 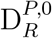. We obtain the rotational diffusion coefficient of the DPPC bilayer for HA free case, 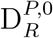 is 0.82 rad^2^/ns. Figure 4(c) shows that 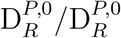 decreases with n_HA5_. The reduced rotational diffusion of the PN vector may indicate the increased electrostatic interactions between HA and DPPC^42^. However, unlike the other quantities, 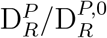 in Figure 4(d) increases linearly with N.

## Discussions

Figure 2(c) and 2(g) show that both the translation and rotation of the water molecules become slower with increasing HA monomer concentration. This can be attributed to the propensity of HA molecules to form hydrogen bonds with water. Our previous study shows that the tetragonal order parameter of water molecules reduces with HA concentration.^8^ The water molecules cannot undergo ordering within themselves due to their increasingly sluggish dynamics around the HA chains. Hence, the water dipoles orient more randomly with more and more HA monomers accessible. The slower translational and rotational motion of the water molecules with HA concentration is in general agreement with the water dynamics in the vicinity of HA and other hydrophilic molecules like dextran^11,12^. The previous report on slowing down of aqueous dynamics in presence of hyaluronidase^17^ is qualitatively similar to the hydrophilic interaction induced slowing down that we observe. Additionally, the HA rotation dynamics, as shown in Figure 3(e), become slower with monomer concentration due to less availability of space for movement. On the other hand, the lipid headgroups do not show direct coupling with HA. However, as HA concentration increases the in-plane diffusion of the lipid head groups increases (Figure 4(a)). The enhanced diffusivity with n_HA5_ is consistent with increased membrane flexibility, reported previously^17^. We note that the lipid is accompanied by slower rotational diffusion of the PN vectors with n_HA5_ (Figure 4(c)), which may be attributed to higher electrostatic interactions^42^.

More significantly, the N dependence of the interfacial dynamics is subtle compared to those for n_HA5_. Water diffusion changes only marginally with N (Figure 2(d) and 2(h)). The HA chain rotational diffusion in 3(f) is also independent of N for larger N. Similarly, the change in lipid lateral diffusion with N is only marginal (Figure 4(b)). The subtle N dependence of the interfacial water dynamics in is quite similar to the polyacrylamide solutions, where water orientational relaxation is found to be chain length independent^43^. As polyacrylamide is a hydrophilic molecule and forms hydrogels similar to HA, the marginal N-dependent dynamical characteristics might be typical of hydrophilic molecules.

To find out the origin of the distinct concentration and chain size dependence of the water dynamics, we consider the organization of the HA chains via the cross-chain radial distribution^44^ of HA monomers, g_xy_(r), where r is the distance in the plane parallel to the bilayer, between the COM of two HA monomers belonging to two distinct chains. Figure 5(a) shows g^xy^(r) vs r plot for different n_HA5_. The cross-chain correlation enhances with n_HA5_. Figure 5(b) shows g^xy^(r) vs r plot for different N. g^xy^(r) has well defined first peak for N=1. The peak disappears with increasing N, indicating that the inter-chain correlations decrease as N increases.

**Figure 5.**
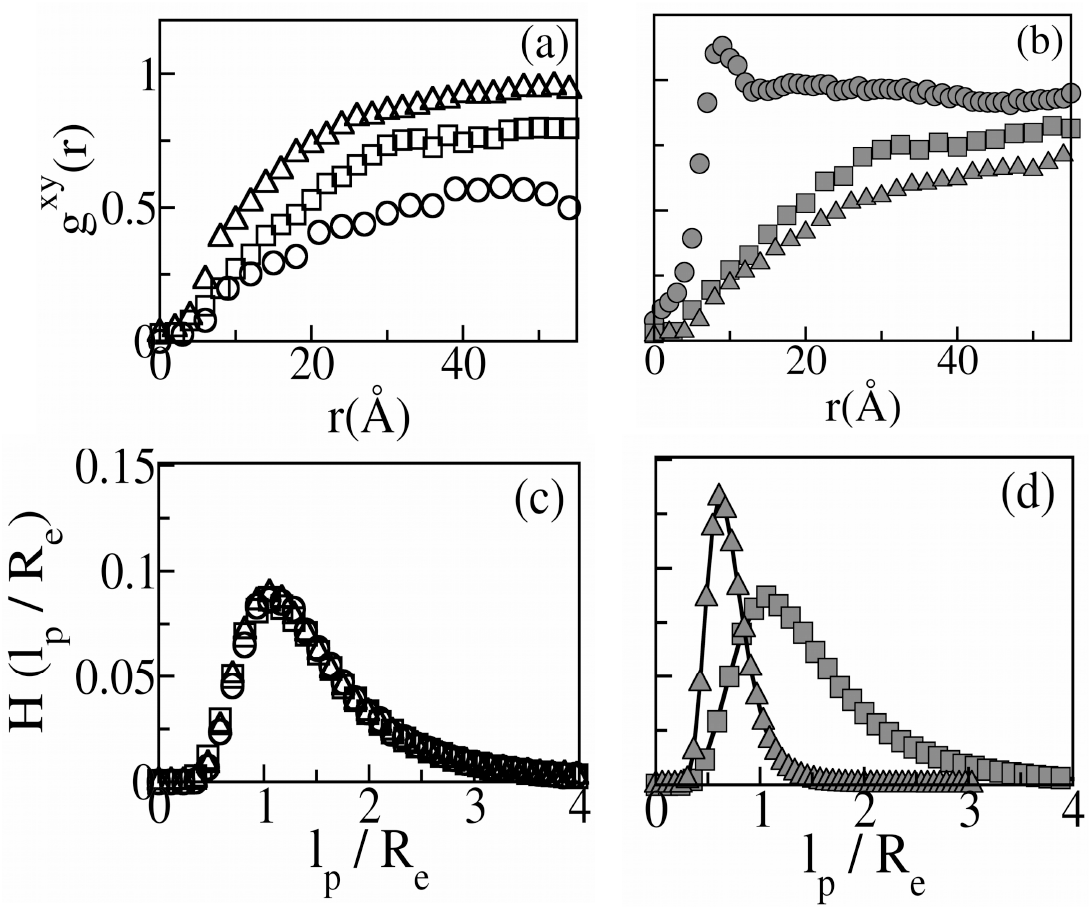
(a) Cross-chain radial distribution of HA monomers at the interface g_xy_(r), in the plane parallel to the bilayer surface, for n_HA5_=10 (*circles*), n_HA5_=30 (*squares*), and n_HA5_=50 (*diamonds*) (open symbols) and (b) for N=1 (*circles*), N=5 (*squares*) and N=10 (*diamonds*) (closed symbols). (c)Histogram of persistence length (*l*_*P*_) to end-to-end distance (R_*e*_) ratio of the HA chains for different n_HA5_ and (d) for different N. Same symbols as (a) and (b) are used.

We also compute the ratio of the persistence length^45^ of the HA chains (l_*P*_) to the end-to-end distances (R_*e*_). l_*P*_ describes the length scale of correlation between the monomers, higher l_*P*_ suggesting a more rigid polymer chain^46^. R_*e*_ is given by the magnitude of the vector joining the COM of the terminal monomers. We show the distribution of l_*P*_ /R_*e*_, H(l_*P*_ /R_*e*_) in Figure 5(c) and Figure 5(d) for different n_HA5_ and N respectively. The peak position of the distribution is the same for different n_HA5_. This indicates that the flexibility of the HA chains remains almost the same for different HA concentrations. Whereas, as we increase N, it shifts to a smaller value of l_*P*_ /R_*e*_. Therefore, the flexibility of the HA molecules increases as HA chain size increases, which in turn weakens the inter-chain correlations due to greater chain flexibility in Figure 5(b). In polymer gels, the diffusivity of water molecules^47^, polystyrene tracer beads^48^ etc. increases with a decrease in network pore size relative to the particle size. Hence, for varying HA chain sizes, the interfacial water molecules experience competition between the hydrophilic interaction with HA and HA chain flexibility. This leads to the dynamics with marginal N dependence. However, the PN rotation becomes faster with N. As we keep the same monomer concentration here, the faster PN rotation may be the result of the higher flexibility of the larger HA chains at the interface.

## Conclusions

To summarize, we study the dynamics of water, short HA chains, and lipids at the interface of HA-water and DPPC bilayer. We observe that the increasing monomer concentration slows down the interfacial dynamics, but the chain size dependence is only marginal. We explain this behavior in terms of the HA network structure and its flexibility at the interface which we expect to hold for all hydrophilic chains. The dynamics we report here are amenable to various spectroscopic experiments, typically in the terahertz (THz) range. The interfacial dynamics near lipids are important in biomedical applications, such as drug delivery to cells^49^, designing HA-coated liposomes^50–52^and development of HA hydrogels^53^.

## Notes

The authors declare no competing financial interest.

## Acknowledgement

A.P. thanks Technical Research Centre at S N Bose National Centre for Basic Sciences, Kolkata for computational facilities and the Council of Scientific and Industrial Research (CSIR), India for financial support [File No. 09/575(0134)/2020-EMR-I].

