## Supporting Information for "Dynamics of aqueous suspension of short Hyaluronic acid chains near DPPC bilayer"

### Methods

#### System Preparation and Force field details

The initial HA structure rcsb pdb id 2BVK[S1] is modified in CHARMM GUI glycan modeller[S2] to obtain HA of different molecular weight (HA1, HA5 and HA10). The DPPC bilayer is set up using CHARMM GUI membrane builder module[S3] in a simulation box of size 12.45 nm $\times$ 12.45 nm $\times$ 19.1 nm. The bilayer consists of a total of 512 lipid molecules with 256 molecules in each monolayer. It spans in the xy plane and its normal is along the z-axis. Aqueous HA with the DPPC bilayer system is prepared using Packmol[S4]. The potential energy of interaction is modeled using CHARMM36 force-field parameters[S5] and TIP3P water model is used for hydrating the system. This is validated previously in the study of HA-lipid systems in literature[S6–S8].

#### Molecular Dynamics (MD) Simulation

All-atom molecular dynamics simulations are performed using the GROMACS package[S9]. Sodium ions are added to each system to maintain electroneutrality since HA contains negatively charged COO- groups. 150 millimolars of Sodium-chloride salt is added to the system to keep the physiological salt concentration. The cut-off lengths for Lennard-Jones and short-range electrostatic interactions are set to 12 Å, while the Particle Mesh Ewald method is used to assess long-ranged electrostatic energy. The HA-free DPPC lipid bilayer system in water is energy minimized using the steepest descent algorithm and followed by short canonical (NVT) and isothermal-isobaric (NPT) equilibrations in water with gradually removing the position and dihedral restraints. The temperature is fixed at 323K using Nose-Hoover thermostat[S10] and pressure is fixed at 1 bar using semi-isotropic pressure coupling with Parrinello-Rahman barostat[S11]. Finally, a production run in NPT ensemble is performed for 10 nanoseconds. The equilibrated final structure of the bilayer is then taken and HA molecules are randomly placed above the bilayer. The HA-DPPC systems, after energy minimization, are again equilibrated through NVT and NPT simulations using position restraints on heavy atoms. The LINCS algorithm is used to constrain all covalent bonds involving hydrogen atoms during the production runs. The final production runs are executed for 600 nanoseconds (ns) in NPT ensemble with 1 femtosecond(fs) integration time step using Leap-Frog integrator removing all restraints and with periodic boundary conditions in all directions. The static results are acquired from the last 400ns trajectory whereas the dynamics data are averaged over three MD trajectories starting from different equilibrium configurations and stored at the time interval of 0.5ps. VMD software[S12] is used to visualize simulation snapshots.

#### Analysis

##### Density Profile Calculations

The density profile of a chemical constituent $\alpha$, $\rho_\alpha(z)$ is defined as the density of $\alpha$ present per unit volume within the thin slices of the same cross-sectional area in the XY plane. Here $\alpha =$W,$\alpha =$H, $\alpha =$P stands for water, the center of mass (COM) of the HA monomers, and phosphorus atoms of the bilayer respectively. We compute $\rho_\alpha(z)$ using the gmx density tool. $\rho_\alpha(z)$ is computed from four distinct blocks, each of 100ns length, extracted from equilibrated trajectories. The error bar is determined by dividing the standard deviation of $\rho_\alpha(z)$ from all the blocks by the square root of the number of blocks[S13].

##### Residence Time Calculations

We define a function, $P_i(t)$, such that $P_i(t) = 1$ if the z-coordinate of the i-th molecule of a given species stays at the interface and $P_i(t) = 0$, if not. The survival probability of species $\alpha$ is defined by the auto-correlation function of $P_i(t)$ given by[S14]

$$S^\alpha = \sum_{i=1}^{N_\alpha} \langle \Pi_{t_k=t_0}^{t_0+t} P_i(t_k) \rangle \tag{1}$$

Here $N_\alpha$ is the total number of species in the system. Angular bracket implies average over multiple time origin $t_0$. We fit a bi-exponential function of the form $ae^{-t/\tau_1} + (1-a)e^{-t/\tau_2}$ to the survival probability and obtain the average residence time as $\tau_\alpha = a\tau_1 + (1-a)\tau_2$.

##### Translational and Rotational Mean Squared Displacements

*Translational MSD*:

The translational mean squared displacements (MSD) of a chemical moiety $\alpha$ at the HA-water and DPPC interface along the bilayer plane (xy plane in our case) are calculated using the formula:

$$< r_\alpha^2(t) >= \frac{1}{N_\alpha} \sum_{i=1}^{N_\alpha} < (x(t) - x(0))^2 + (y(t) - y(0))^2 > \tag{2}$$

Here $\alpha$ =W,$\alpha$ =H, $\alpha$ =P stands for water, HA monomers, and phosphorus atoms of the bilayer respectively. $< r_\alpha^2 >$ is computed over only those molecules which remain at the interface continuously for time t[S15]. Their number is indicated by $N_\alpha$ here. The angular bracket denotes the average over multiple time origins. While computing the lateral mean squared displacement of the phosphorus atoms of the lipid bilayer, the COM motion of the respective leaflet is removed for each lipid molecule[S16,S17].

*Rotational MSD*:

The rotational MSD $\langle \phi_\alpha^2 \rangle$ of $\alpha$, is computed using the following definition[S14,S18]

$$\langle \phi_\alpha^2(t) \rangle = \frac{1}{N_\alpha} \sum_{i=0}^{N_\alpha} \langle |\vec{\phi}_i(t_0 + t) - \vec{\phi}_i(t_0)|^2 \rangle \tag{3}$$

Here $t_0$ is an arbitrary time origin and the vector rotational displacement $\vec{\phi}_i(t_0 + t)$ is the sum of infinitesimal angular displacements $\delta\vec{\phi}_i$ of a suitably chosen vector associated with the molecule from $\vec{\phi}_i(t_0)$ to $\vec{\phi}_i(t_0+t)$ in discrete time steps $\delta t$. Mathematically, $\vec{\phi}_i(t_0+t) - \vec{\phi}_i(t_0) = \int_{t_0}^{t_0+t} \delta\vec{\phi}_i(t')$[S18,S19]. For water and HA $[\vec{\phi}_i(t) \equiv \vec{\phi}_w(t)$ and $\vec{\phi}_i(t) \equiv \vec{\phi}_h(t)]$ is calculated from the water dipole vector $\vec{\mu_i^w}$ and HA end-to-end vectors. For lipids, $\vec{\phi}_{pn}(t)$ is calculated using the projection of the P-N vector in the xy plane[S20].

For water, the magnitude of $\delta\vec{\phi}_w$ is $cos^{-1}[\hat{\mu_i^w}(t).\hat{\mu_i^w}(t+\delta t)]$ and its direction is along $\hat{\mu_i^w}(t) \times \hat{\mu_i^w}(t+\delta t)$. Note that any suitably chosen vector other than $\mu_i^w$ can be used to describe the rotational diffusion, but they yield the same qualitative conclusion[S18,S21,S22]. For HA and PN vectors, we follow the same method as well. With this technique $\vec{\phi_i}(t)$ becomes an unbounded variable. $N_\alpha$ in equation 3 indicates the number of $\alpha$ species at the interface throughout the time interval of t. $\langle \phi_\alpha^2 \rangle$ is fitted with a straight line to acquire the rotational diffusion $D_R^\alpha$ of $\alpha$ at the interface[S14].

$$D_R^\alpha = \lim_{t\to\infty} \frac{1}{4t} \langle \phi_\alpha^2 \rangle \tag{4}$$

For waters, we also compute rotational correlation time $\tau_l^W$ from rotational autocorrelation function[S15]. We verified that the product of $\tau_l^W$ and $D_R^W$ remains constant[S23].

The error bars for the residence times and diffusion coefficients are derived from the standard deviation of the respective quantities from three separate MD runs divided by the square root of the number of runs[S13]. While scaling the data by HA-free cases (see Results), we compute the error bars using the error propagation method of division[S24].

#### Orientational autocorrelation function

The orientational autocorrelation function ($C(\tau)$) is computed over only those water molecules that continuously stay at the interface. $C(\tau)$ is defined as follows :

$$C(\tau) = \langle \frac{\vec{\mu_i}(\tau).\vec{\mu_i}(\tau)}{\vec{\mu_i}(0).\vec{\mu_i}(0)} \rangle \tag{5}$$

where $\vec{\mu_i}(\tau)$, the dipole vector of the ith water molecule at time $\tau$, is defined as the vector pointing from the oxygen atom to the center of mass of two hydrogen atoms[S15]. C($\tau$) is fitted with a bi-exponential function of the form $C(\tau) = a_1 e^{-\tau/t1} + (1-a_1)e^{-\tau/t2} + b$ to obtain different timescales. The parameter b is used to take care of the long-lived tail of the C($\tau$). The average orientation timescale is obtained by $\tau^{\mathrm{w}} = a_1 t_1 + (1-a_1)t_2$.

#### Cross-chain Radial Distribution

We compute cross-chain radial distribution function $g_{xy}$(r) of the HA monomers the parallel plane to the bilayer (xy plane) using the following formula[S25]:

$$g_{xy}(r) = \langle \frac{1}{\rho_N} \sum_{i=1}^{N} \sum_{j=1}^{N} \delta(|\vec{r_i} - \vec{r_j}|_{xy} - r) \rangle \tag{6}$$

Here $\vec{r_i}$ and $\vec{r_j}$ indicates position vector of the com of two HA monomers belonging to different chains at the interface, $r = \sqrt{x^2 + y^2}$, $|\vec{r_i} - \vec{r_j}|_{xy} = \sqrt{(x_i - x_j)^2 + (y_i - y_j)^2}$ and $\rho_N$ implies the areal monomer density (in xy plane) at the interface. N is the number of beads at the interface. Angular bracket represents average over snapshots.

#### Persistence Length Calculations

We calculate the persistence length of HA ($l_p$) the following formula[S26] :

$$l_p = \frac{-b}{ln < cos\theta >}$$

Here b is the average distance between the center of masses of the HA monomers. $\theta$ is the angle between the center of masses of three consecutive HA monomers. Note that $l_p$ is not defined for HA monomers (N=1).

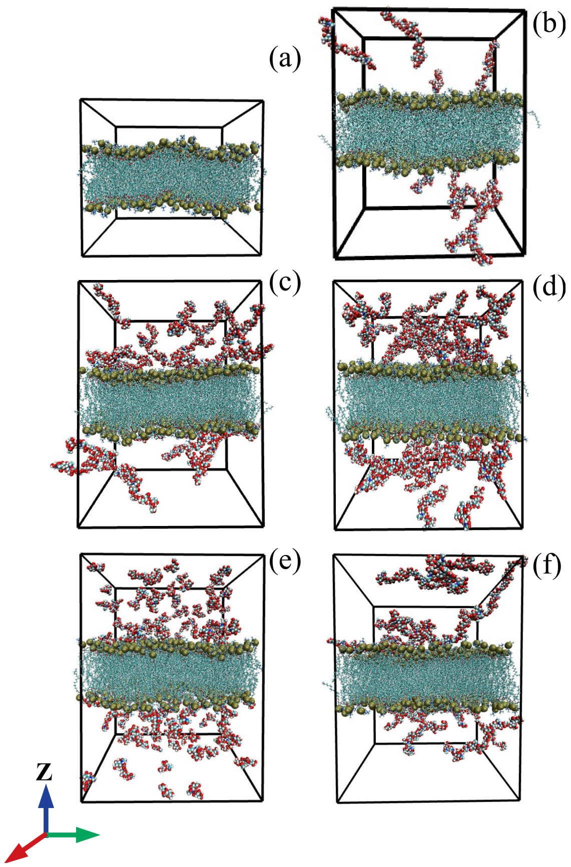


**Figure S1.** Typical equilibrium snapshot of (a) $n_{HA5}$=0, (b) $n_{HA5}$=10, (c) $n_{HA5}$=30, (d) $n_{HA5}$=50, (e) N=1 and (f) N=10. Hyaluronic acids (HA) and lipid bilayer are shown in vdw and bonds representation respectively with the following color codes: *Cyan* for Carbons, *red* for Oxygens, and *blue* for Nitrogens. The normal of the lipid bilayer is along z axis. Water molecules are not shown.

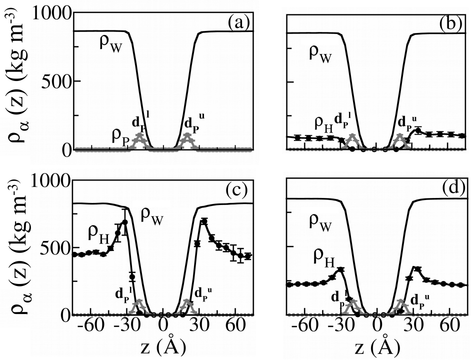


**Figure S2.** (a) Density profiles of Phosphorus atoms ($\rho_P$(z), *grey solid line*), HA ($\rho_H$(z), *dotted black line*), and water ($\rho_W$(z), *solid black line*) along the bilayer normal for $n_{HA5}$=0, (b) $n_{HA5}$=10, (c) $n_{HA5}$=50 and (d) N=1. Origin is set at the bilayer center. ${d_P}^l$ and ${d_P}^u$ show the peaks of $\rho_P$(z). $\rho_H$(z) is amplified by a factor of 5.

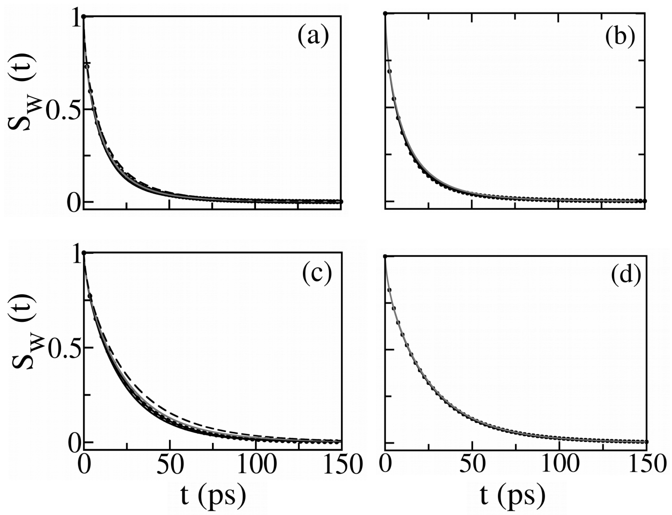


**Figure S3.** (a) Survival probability of water $S_W(t)$ in region A for different HA concentrations: $n_{HA5}$=0 (*solid black line*), $n_{HA5}$=10 (*dotted black line*), $n_{HA5}$=30 (*solid gray line*), $n_{HA5}$=50 (*dashed black line*) and (b) for varying HA chain size: N=1 (solid black line), N=1 (dotted black line), N=10 (solid gray line). (c) $S_W(t)$ in region B for different $n_{HA5}$ and (d) for different N. The same line type as (a)-(b) is used.

**Table S1.** Mean residence time of water in region A ($\tau_W^A$) and in region B ($\tau_W^B$), obtained from $S_W(t)$ in respective regions (Figure S3). $\tau_W^{A,0}$ and $\tau_W^{B,0}$ are shown.

| system | $\tau_W^A$ (ps) | $\tau_W^B$ (ps) |
|---|---|---|
| $n_{HA5}$=0 | 10.86 ($\tau_W^{A,0}$) | 20.07 ($\tau_W^{B,0}$) |
| $n_{HA5}$=10 | 11.86 | 21.24 |
| $n_{HA5}$=30 | 11.83 | 22.90 |
| $n_{HA5}$=50 | 13.40 | 25.27 |
| N=1 | 12.37 | 22.32 |
| N=5 | 11.83 | 22.90 |
| N=10 | 12.79 | 23.08 |

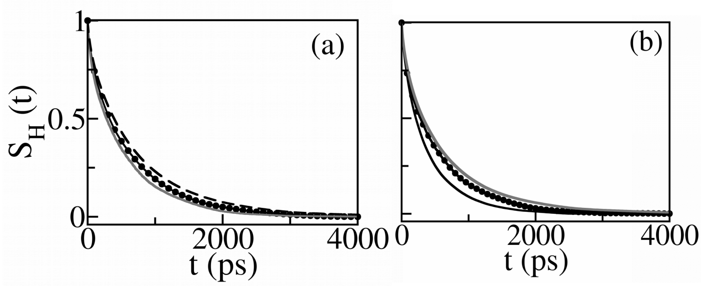


**Figure S4.** Survival probability of HA monomers $S_H(t)$ in region B for (a) $n_{HA5}$=10 (*dotted black line*), $n_{HA5}$=30 (*solid gray line*), $n_{HA5}$=50 (*dashed black line*), and (b) N=1 (*solid black line*), N=5 (*dotted black line*), N=10 (*solid gray line*).

**Table S2.** Mean residence time of HA monomers ($\tau_H$) in region B, computed from $S_H(t)$ (Figure S4). $\tau_H^1$ is indicated.

| system | $\tau_H$ (ps) |
|---|---|
| $n_{HA5}$=0 | - |
| $n_{HA5}$=10 | 568.84 |
| $n_{HA5}$=30 | 494.87 |
| $n_{HA5}$=50 | 677.04 |
| N=1 | 372.48 ($\tau_H^1$) |
| N=5 | 494.87 |
| N=10 | 588.65 |

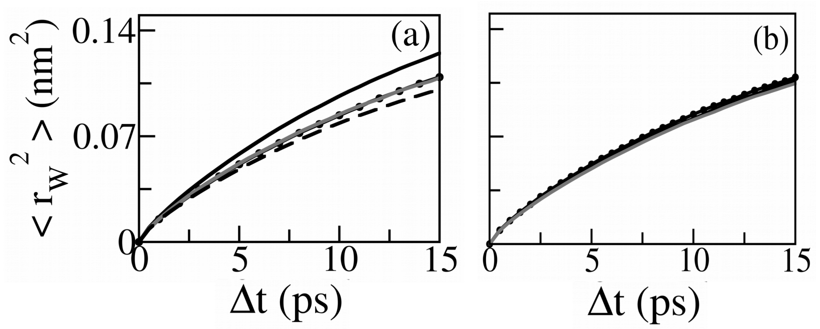


**Figure S5.** Translational MSD of the water molecules ($\langle r_W^2 \rangle$) in region A, in the plane parallel to the bilayer surface, for different HA concentrations: $n_{HA5}$=0 (*solid black line*), 10 (*dotted black line*), 30 (*solid gray line*) and 50 (*dashed black line*) (b) $\langle r_W^2 \rangle$ in region B for varying HA chain sizes: N=1 (*solid black line*), N=5 (*dotted black line*) and N=10 (*solid gray line*)

**Table S3.** Translational diffusion exponents ($\beta_W$) of $\langle r_W^2 \rangle$ in region A

| system | $\beta_W$ |
|---|---|
| $n_{HA5}$=0 | 0.75 |
| $n_{HA5}$=10 | 0.73 |
| $n_{HA5}$=30 | 0.73 |
| $n_{HA5}$=50 | 0.72 |
| N=1 | 0.73 |
| N=5 | 0.73 |
| N=10 | 0.73 |

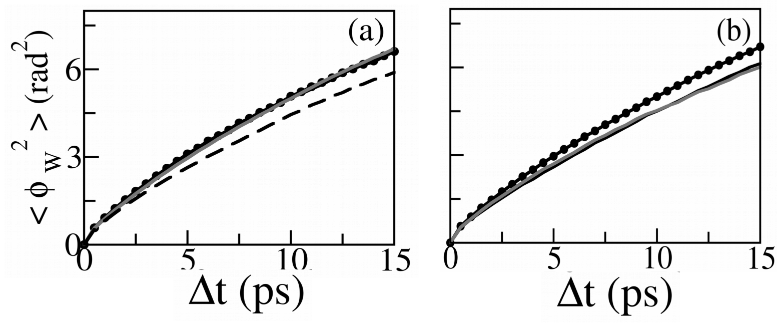


**Figure S6.** Rotational MSD of the water molecules ($\langle \phi_W^2 \rangle$) in region A for different HA concentrations: $n_{HA5}$=0 (*solid black line*), 10 (*dotted black line*), 30 (*solid gray line*) and 50 (*dashed black line*) (b) $\langle r_W^2 \rangle$ in region B for varying HA chain sizes: N=1 (*solid black line*), N=5 (*dotted black line*) and N=10 (*solid gray line*)

**Table S4.** Rotational diffusion exponents ($\gamma_W$) of $\langle \phi_W^2 \rangle$ in region A

| system | $\gamma_W$ |
|---|---|
| $n_{HA5}$=0 | 0.73 |
| $n_{HA5}$=10 | 0.72 |
| $n_{HA5}$=30 | 0.74 |
| $n_{HA5}$=50 | 0.72 |
| N=1 | 0.73 |
| N=5 | 0.73 |
| N=10 | 0.75 |

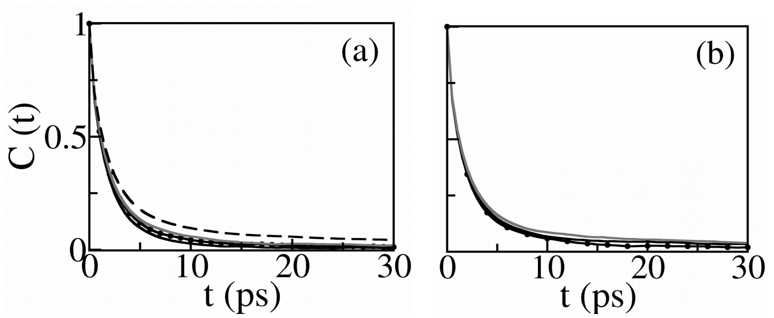


**Figure S7.** Rotational autocorrelation function of the water molecules (C($\tau$)) in region B for different HA concentrations: $n_{HA5}$=0 (*solid black line*), 10 (*dotted black line*), 30 (*solid gray line*) and 50 (*dashed black line*) (b) $\langle r_W^2 \rangle$ in region B for varying HA chain sizes: N=1 (*solid black line*), N=5 (*dotted black line*) and N=10 (*solid gray line*)

**Table S5.** Rotational autocorrelation time of water in region B, $t_1^W$ (computed from C(t) in SI Figure S7) and its product with rotational diffusion, $D_R^W$ (for first rank, here l=1)

| system | $t_1^W$(ps) | $t_1^W \cdot D_R^W$ |
|---|---|---|
| $n_{HA5}$=0 | 1.84 | 0.51 |
| $n_{HA5}$=10 | 2.13 | 0.55 |
| $n_{HA5}$= 30 | 2.21 | 0.56 |
| $n_{HA5}$= 50 | 2.53 | 0.59 |
| N=1 | 2.13 | 0.53 |
| N=5 | 2.21 | 0.56 |
| N=10 | 2.53 | 0.60 |

**Table S6.** translational diffusion exponents ($\beta_H$) of HA monomers in region B.

| system | $\beta_H$ |
|---|---|
| $n_{HA5}$=0 | - |
| $n_{HA5}$=10 | 0.80 |
| $n_{HA5}$= 30 | 0.80 |
| $n_{HA5}$= 50 | 0.79 |
| N=1 | 0.88 |
| N=5 | 0.80 |
| N=10 | 0.79 |

**Table S7.** First rank rotational autocorrelation time $t_1^H$ for interfacial HA monomers

| system | $t_1^H$ (ns) |
|---|---|
| $n_{HA5}$=0 | - |
| $n_{HA5}$=10 | 6.1 |
| $n_{HA5}$= 30 | 6.8 |
| $n_{HA5}$= 50 | 7.7 |
| N=1 | 0.2 |
| N=5 | 6.8 |
| N=10 | 24.6 |

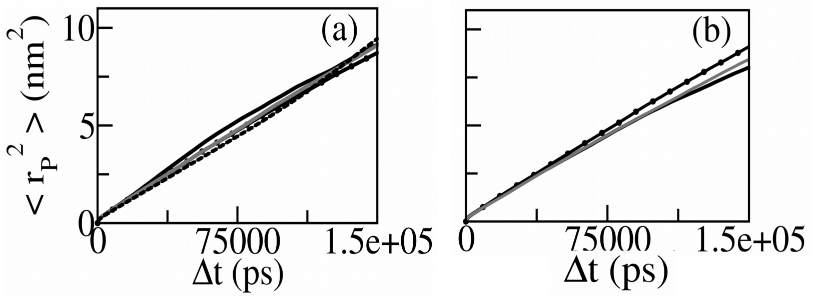


**Figure S8.** (a) Translational mean squared displacements of the Phosphorus atoms ($\langle r_P^2 \rangle$) in the lateral plane of the bilayer for varying HA concentrations: $n_{HA5}$ = 0 (*solid black line*), 10 (*dotted black line*), 30 (*gray solid line*), 50 (*dashed black line*) and (b) for different HA chain sizes: N=1 (*solid black line*), N=5 (*dotted black line*) and N=10 (*solid gray line*)

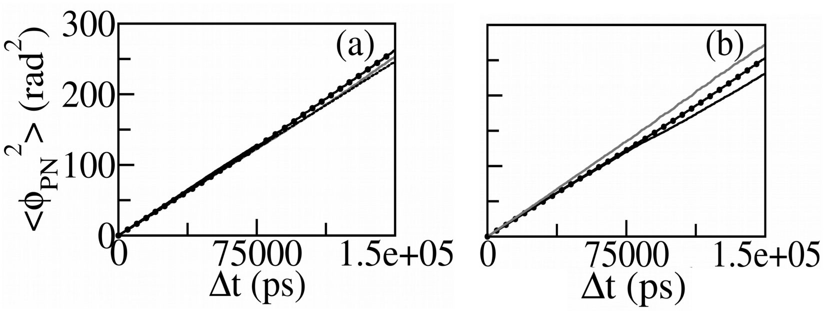


**Figure S9.** (a) Rotational mean squared displacements of the lipid PN vectors ($\langle \phi_{PN}^2 \rangle$) for $n_{HA5}$ = 0 (*solid black line*), 10 (*dotted black line*), 30 (*gray solid line*), 50 (*dashed black line*) and (b) for N=1 (*solid black line*), N=5 (*dotted black line*) and N=10 (*solid gray line*)

[S3] Wu, E. L.; Cheng, X.; Jo, S.; Rui, H.; Song, K. C.; Dávila-Contreras, E. M.; Qi, Y.; Lee, J.; Monje-Galvan, V.; Venable, R. M.; others CHARMM-GUI membrane builder toward realistic biological membrane simulations. 2014.

[S4] Martínez, L.; Andrade, R.; Birgin, E. G.; Martínez, J. M. PACKMOL: A package for building initial configurations for molecular dynamics simulations. *Journal of computational chemistry* **2009**, *30*, 2157–2164.

[S5] Guvench, O.; Mallajosyula, S. S.; Raman, E. P.; Hatcher, E.; Vanommeslaeghe, K.; Foster, T. J.; Jamison, F. W.; MacKerell Jr, A. D. CHARMM additive all-atom force field for carbohydrate derivatives and its utility in polysaccharide and carbohydrate–protein modeling. *Journal of chemical theory and computation* **2011**, *7*, 3162–3180.

[S6] Smith, P.; Ziolek, R. M.; Gazzarrini, E.; Owen, D. M.; Lorenz, C. D. On the interaction of hyaluronic acid with synovial fluid lipid membranes. *Physical Chemistry Chemical Physics* **2019**, *21*, 9845–9857.

[S7] Paul, D.; Paul, A.; Mukherjee, D.; Saroj, S.; Ghosal, M.; Pal, S.; Senapati, D.; Chakrabarti, J.; Pal, S. K.; Rakshit, T. A Mechanoelastic Glimpse on Hyaluronan-Coated Extracellular Vesicles. *The Journal of Physical Chemistry Letters* **2022**, *13*, 8564–8572.

[S8] Zhang, H.; Cai, W.; Shao, X. Regulation of aquaporin-3 water permeability by hyaluronan. *Physical Chemistry Chemical Physics* **2021**, *23*, 25706–25711.

[S9] Van Der Spoel, D.; Lindahl, E.; Hess, B.; Groenhof, G.; Mark, A. E.; Berendsen, H. J. GROMACS: fast, flexible, and free. *Journal of computational chemistry* **2005**, *26*, 1701–1718.

[S10] Nosé, S. A molecular dynamics method for simulations in the canonical ensemble. *Molecular physics* **1984**, *52*, 255–268.

[S11] Parrinello, M.; Rahman, A. Polymorphic transitions in single crystals: A new molecular dynamics method. *Journal of Applied physics* **1981**, *52*, 7182–7190.

[S12] Humphrey, W.; Dalke, A.; Schulten, K. VMD: visual molecular dynamics. *Journal of molecular graphics* **1996**, *14*, 33–38.

[S13] Venable, R. M.; Brown, F. L.; Pastor, R. W. Mechanical properties of lipid bilayers from molecular dynamics simulation. *Chemistry and physics of lipids* **2015**, *192*, 60–74.

[S14] Debnath, A.; Mukherjee, B.; Ayappa, K.; Maiti, P. K.; Lin, S.-T. Entropy and dynamics of water in hydration layers of a bilayer. *The Journal of Chemical Physics* **2010**, *133*, 174704.

[S15] Hande, V. R.; Chakrabarty, S. How far is "bulk water" from interfaces? Depends on the nature of the surface and what we measure. *The Journal of Physical Chemistry B* **2022**, *126*, 1125–1135.

[S16] Siu, S. W.; Vácha, R.; Jungwirth, P.; Böckmann, R. A. Biomolecular simulations of membranes: physical properties from different force fields. *The Journal of chemical physics* **2008**, *128*.

[S17] Venable, R. M.; Ingólfsson, H. I.; Lerner, M. G.; Perrin Jr, B. S.; Camley, B. A.; Marrink, S. J.; Brown, F. L.; Pastor, R. W. Lipid and Peptide Diffusion in Bilayers: The

Saffman–Delbruck Model and Periodic Boundary Conditions. *The Journal of Physical Chemistry B* **2017**, *121*, 3443–3457.

[S18] Mazza, M. G.; Giovambattista, N.; Starr, F. W.; Stanley, H. E. Relation between rotational and translational dynamic heterogeneities in water. *Physical review letters* **2006**, *96*, 057803.

[S19] Tendong, E.; Dasgupta, T. S.; Chakrabarti, J. Dynamics of water trapped in transition metal oxide-graphene nano-confinement. *Journal of Physics: Condensed Matter* **2020**, *32*, 325101.

[S20] Moore, P. B.; Lopez, C. F.; Klein, M. L. Dynamical properties of a hydrated lipid bilayer from a multinanosecond molecular dynamics simulation. *Biophysical journal* **2001**, *81*, 2484–2494.

[S21] Mazza, M. G.; Giovambattista, N.; Stanley, H. E.; Starr, F. W. Connection of translational and rotational dynamical heterogeneities with the breakdown of the Stokes-Einstein and Stokes-Einstein-Debye relations in water. *Physical Review E* **2007**, *76*, 031203.

[S22] Zhang, Q.; Wu, T.; Chen, C.; Mukamel, S.; Zhuang, W. Molecular mechanism of water reorientational slowing down in concentrated ionic solutions. *Proceedings of the National Academy of Sciences* **2017**, *114*, 10023–10028.

[S23] Lombardo, T. G.; Debenedetti, P. G.; Stillinger, F. H. Computational probes of molecular motion in the Lewis-Wahnström model for ortho-terphenyl. *The Journal of chemical physics* **2006**, *125*, 174507.

[S24] https://www.geol.lsu.edu/jlorenzo/geophysics/uncertainties/Uncertaintiespart2.html.

[S25] Karmakar, R.; Ghosal, A.; Chakrabarti, J. Model studies on motion of respiratory droplets driven through a face mask. *Europhysics Letters* **2022**,

[S26] Nikoubashman, A.; Milchev, A.; Binder, K. Dynamics of single semiflexible polymers in dilute solution. *The Journal of Chemical Physics* **2016**, *145*, 234903.
